# PRIME-BSPre: A genome-wide protein-RNA binding sites prediction method based on templates

**DOI:** 10.1101/2023.10.08.561403

**Authors:** Xinhang Wei, Yingtian Duan, Danyang Li, Xudong Liu, Juan Xie, Shiyong Liu

## Abstract

Identification of RNA binding sites that potentially interact with RNA-binding proteins facilitates a comprehensive analysis of protein-RNA interactions and enables further investigation into the mechanisms underlying RNA splicing and modification. However, the current experimental data remains limited in comparison to the vast family of RBPs, and deep learning prediction methods are inadequate for those RBPs lacking sufficient interaction data for training. Therefore, we present PRIME-BSPre, a genome-wide method for predicting protein-RNA binding sites based on templates that incorporate both RNA sequence and secondary structure as well as the tertiary structure of corresponding RBPs. We have successfully benchmarked our method on the human genome, demonstrating excellent prediction performance on RBP datasets beyond our library and robustness across cell lines. Additionally, we are pioneers in introducing the low Shannon entropy algorithm to describe binding preferences of RNA motifs. Our predicted results further support the hypothesis that RBPs preferentially bind RNA motifs with low complexity.

## INTRODUCTION

Post-transcriptional regulation is a crucial aspect of gene expression, and RNA-binding proteins (RBPs) play key roles in this process (Spitale et al. 2015b; England et al. 2022). To fully comprehend the physiological functions and potential contributions of RBPs, it is imperative to identify their interacting RNAs and specific binding sites(Van Nostrand et al. 2020). Enhanced CLIP (eCLIP) is an in vivo method derived from CLIP-seq (crosslinking and immunoprecipitation followed by sequencing), which enables the study of protein-RNA interaction on a genomic scale(Van Nostrand et al. 2016). The eCLIP output data have demonstrated utility in investigating the functional roles and binding preferences of RBPs (Gerstberger et al. 2014; Van Nostrand et al. 2020), and are widely used to predict the possible protein-RNA interactions (Laverty et al. 2022; Xu et al. 2023). Compared to iCLIP (Konig et al. 2010) and PAR-CLIP (Hafner et al. 2010), eCLIP offers the advantages of simplicity, high success rate, high resolution, and partially addresses the issue of biased base composition of RNA sequences caused by cellular factors. And in comparison, classical in vitro method SELEX (Cook et al. 2015) or, RNAcompete (Cook et al. 2017) and RNA Bind-n-Seq (Lambert et al. 2014) which were developed subsequently, eCLIP is more physiologically relevant and better restores the native structure of RNAs due to its cellular context. Additionally, the nucleotide sequences utilized in eCLIP are more comprehensive and representative of their natural state compared to synthetic sequences used in vitro that were fragmented prior to interacting with RBPs.

Faced with a large number of RBPs and their corresponding RNAs, experimental research alone has its limitation. Therefore, based on the sequencing results obtained through in vivo or in vitro methods, numerous predictive models for protein-RNA interactions have been proposed. Due to the abundance of available data, most prediction methods rely on deep learning models or machine learning techniques such as support vector machines (SVM) (Kazan et al. 2010; Maticzka et al. 2014; Spitale et al. 2015a; Orenstein et al. 2016; Xu et al. 2023). However, training these models requires high-confidence, low-noise and well-structured data. Therefore, researchers prefer to train their models using RNAcompete and RNA Bind-n-Seq datasets rather than CLIP-seq. And this may face the following problems: (1) As previously stated, the in vitro experimental methods utilize synthetic RNA sequence fragments that lack complete secondary structure during binding, thus preventing the acquisition of structural preferences from the data. So certain methods such as RNAcontext (Kazan et al. 2010) or PrismNet (Xu et al. 2023) choose to use prediction methods such as Sfold (Ding and Lawrence 2003) or experimental data obtained from icSHAPE (Spitale et al. 2015a) to supplement the absence of RNA secondary structure in their datasets. However, the supplemented data cannot fully capture the actual binding process, potentially resulting in a discrepancy between sequence-based and structure-based binding preferences learned by models. (2) The binding preference obtained by the aforementioned experimental methods is heavily reliant on RBPs. In other words, each deep learning model can only be trained and accurately run prediction under one specific RBP.

Moreover, investigations into the binding preferences of RBPs have revealed a remarkable conservation in both their RNA binding domains and the specific regions of RNAs they interact with (Ray et al. 2013; Dominguez et al. 2018). Therefore, we aimed to develop a universal prediction model based on these shared characteristics, free from the constraints of specific RBP binding preferences. As such, we have proposed a template-based prediction method PRIME-3D2D in our previous research (Xie et al. 2020). This method can basically complete the prediction on the yeast genome, but its template scoring function only considers the global similarity between RBPs and RNAs, ignoring the local characteristics of binding sites. Therefore, in PRIME-BSPre, we focused on adding local similarity scores to improve predictive performance. Whether utilizing deep learning or classical computational methods, obtaining specific features of the binding sites is crucial for accurately predicting protein-RNA interactions and identifying the strength of their binding affinity. Typically, the features observed at RNA binding sites can be classified into two distinct categories: sequence-based and structure-based. Earlier methods, such as DeepBind (Alipanahi et al. 2015), solely relied on sequence-based information to train models and achieved some success. However, recent approaches like RNAcontext (Kazan et al. 2010) and GraphProt (Hiller et al. 2006; Maticzka et al. 2014) have incorporated RNA secondary structure features into their models, resulting in improved performance. In terms of selecting RNA secondary structure features, most regression models (Laverty et al. 2022) or sequence-based neural network models only consider the pairing information (Orenstein et al. 2016). While some methods serialize the complete secondary structure, a few methods, such as GraphProt, employ graph networks to directly extract both the secondary structural components and their corresponding sequence information.

Experimental evidence suggests that RBPs exhibit a conservative preference for specific RNA motifs, with a particular affinity for low complexity motifs (Dominguez et al. 2018). Additionally, RNA secondary structure is also crucial in protein-RNA interactions (Stefl et al. 2005; Hiller et al. 2006; Dominguez et al. 2018; Sun et al. 2021). Compared to the aforementioned methods that directly learn binding preferences for predicting binding sites, our template-based approach offers the advantage of utilizing native binding sites for the all-to-all preliminary alignment, considering sequence and secondary structure similarities. Besides, tertiary structural similarity between RBPs was also taken into account (Zhang and Skolnick 2005). We propose that when the target and template exhibit high structure and sequence similarity, their binding preferences are likely to be similar as well. And the alignment also revealed that around 6% of the templates exhibited efficacy (MCC > 0.6) in each prediction. Therefore, we further propose a set of scoring and screening approaches based on the sequence and secondary structure of the RNA binding sites to identify effective templates in the all-to-all preliminary alignment. The assessment of binding preference similarity between templates and targets incorporates the complexity of RNA motifs by calculating the Shannon entropy of RNA sequences within an 8-mer sliding window. Moreover, we have devised a novel LS-2d approach for quantifying secondary structures, enabling the comparison of characteristic structural features between templates and targets (Figure 1).

**Figure 1.**
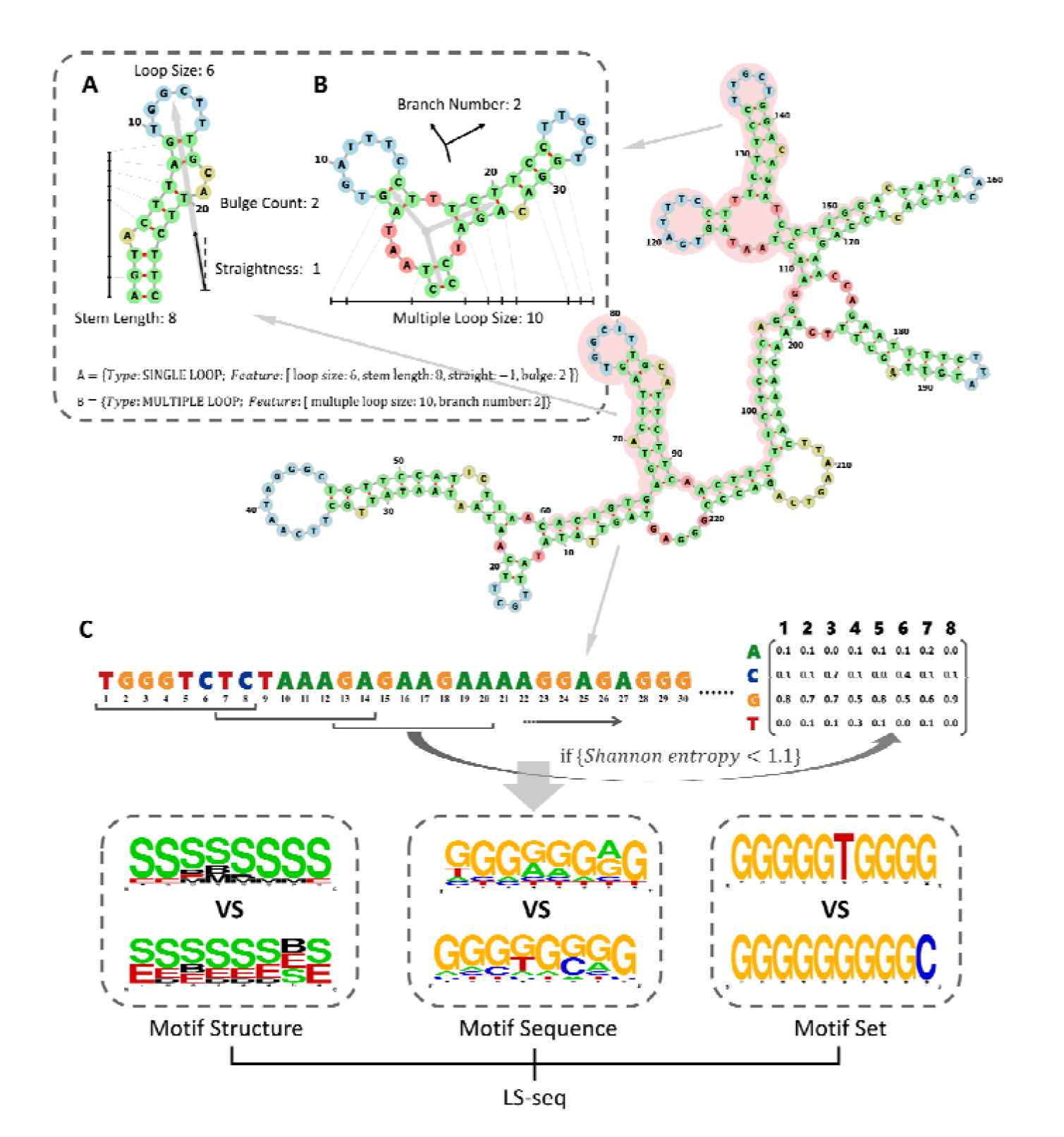
Sequence feature extraction and similarity quantification comparison of RNA binding sites. The pink-shaded region in the RNA secondary structure represents the binding sites, while LS-Score quantitatively assesses sequence and secondary structure similarities within this area between templates and targets. **A** The secondary structure HAIR LOOP, which exhibits typical characteristics, is depicted along with its quantitative features. **B** The secondary structure MULTIPLE LOOP, which exhibits another characteristic feature, is depicted along with its corresponding quantitative attributes. **C** shows the Sequence features of native and predicted RNA binding sites: Motif Structure, Motif Sequence and Motif Set. Among them, Motif Structure is the PPW generated by serialized secondary structure features: B (Bulge), E (External strand), H (Hair loop), M (Multiple loop) and S (Stem); Motif set is a base enriched in motifs.

**Table 1.**
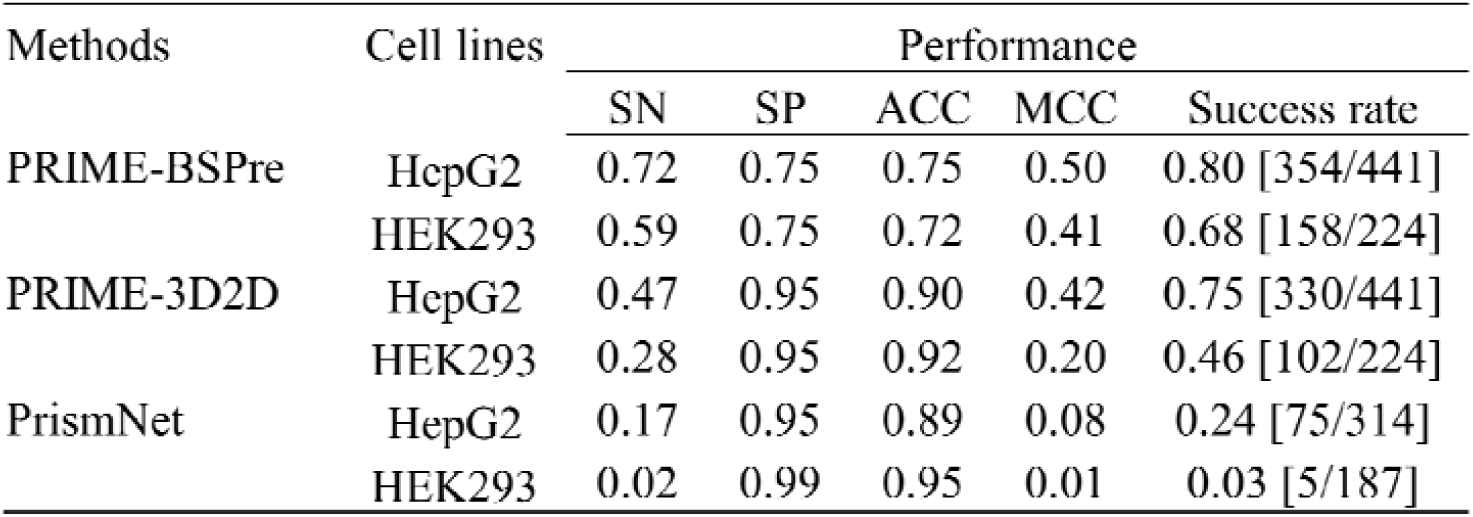
The performance of PRIME-BSPre versus other methods on different cell lines (success defined as MCC>0.1)

## METHODS

### Template library construction

Our template library was constructed using 158 human RBPs, comprising of 15,348 RNA templates that interact with them (see Table S1). In contrast to the studies focusing solely on RNA fragments, our templates accurately map the eCLIP data across the entire genome. All templates were built based on complete transcript sequences. To obtain the RNA binding sites locations of human RBPs, we downloaded eCLIP-bed files from ENCODE (Luo et al. 2020) (https://www.encodeproject.org/) and cross-referenced them with CDS files of human transcripts obtained from Ensembl database (Cunningham et al. 2022) (http://www.ensembl.org/index.html). Using SAMtools (Danecek et al. 2021), we extracted complete transcript sequences from the HUMAN hg38 genome assembly (Lander et al. 2001) (https://hgdownload.soe.ucsc.edu/downloads.html). In addition to considering RNA sequence similarity, our algorithm incorporates the tertiary structures of RBPs and the secondary structures of interacting RNAs. To ensure flexibility and universality in constructing the template library, we utilized RNAstructure (Reuter and Mathews 2010) for predicting RNA secondary structures, albeit with a slight sacrifice in precision. Due to limitations in precision and computational time, the maximum allowed transcript length was restricted to less than 2000nt. Furthermore, we obtained PDB files for 158 RBPs primarily from experimental data available on UniProt (https://www.uniprot.org/). For proteins with incomplete experimental structures, we supplemented them with predicted results from AlphaFold Protein Structure Database (Jumper et al. 2021; Varadi et al. 2022) (https://alphafold.com/).

### All-to-all alignment against the template library

Users need to provide the sequence and secondary structure of the RNA, and the PDB file of their inquired RBP. These inputs are utilized to perform an all-to-all alignment across the entire template library respectively. TMalign (Zhang and Skolnick 2005) is used to align the tertiary structure of the target protein with the RBPs in the template library, yielding a similarity score (TM-Score). Additionally, the sequence and secondary structure of the target RNA are globally aligned with the RNAs in the library using LocARNA (Will et al. 2012). By integrating the sequence-based and structure-based information, LocARNA can generate sequences alignment between the target and template. Utilizing these alignment results allows us to accurately map native binding sites from templates onto the corresponding positions within the target RNA sequence, ultimately providing preliminary prediction outputs.

### A function LS-Score for measuring the local similarity between the native and predicted binding sites

After conducting all-to-all alignment, we observed that approximately 6% of the predictions were deemed high quality (MCC>0.6). However, the original 3D2D-Score (*W* * *TM_score_* + (1 − *W*) * *RNAsimilarity*, *W* = 0.6) (Xie et al. 2020) had a poor screening capability, and only a limited number of effective templates could be identified. 3D2D-Score was constructed for protein-RNA complexes, both TM-Score and RNAsimilarity are global. The 3D2D-score was deemed competent for protein-RNA complex analysis due to the RNA sequences in the complexes are generally short. However, when aiming to identify precise locations of the binding sites within longer RNA sequences, the 3D2D-score alone proved insufficient, necessitating the development of the Local Similarity score (LS-Score). The LS-Score is a scoring function specifically designed for the assessment of similarity between native binding sites on templates and their corresponding predicted binding sites on target RNAs. It comprises two main components, namely LS-2d (local secondary structure similarity) and LS-seq (local sequence binding preference similarity). See Table S2 for specific scoring strategy.

#### LS-2d (Local secondary structure similarity)

We believe that the preliminary alignment may be influenced by strong similarity regions outside the native binding sites, resulting in the actual similarity of the binding sites is not that strong. Thus, we developed LS-2d approach to extract, quantify and compare the characteristic secondary structures between the native and predicted binding sites. We categorized local characteristic secondary structures into two groups: SINGLE LOOP and MULTIPLE LOOP (Figure 1A & Figure 1B). SINGLE LOOP includes loop size, stem length (the number of paired bases in the stem), bulge count (the number of bulges present on the stem), and straightness (by calculating the difference between the size and number of bulges on both sides of the stem, the distribution characteristics of bulges are described). Due to the complexity of MULTIPLE LOOP, our focus was directed towards two key features: branch number and multiple loop size, which were further complemented by a comparative analysis of secondary structures presented in the dot-bracket format. The similarity score is computed separately for each quantified feature, and subsequently integrated into the LS-2d metric. This method systematically explores the characteristic secondary structures of both the target and template, selecting the maximum LS-2d value as the final matching outcome.

#### LS-seq (Local sequence binding preference similarity)

In contrast to the traditional similarity score obtained through sequence alignment, we utilized an 8-mer sliding window with a step size of 6nt to scan both native and predicted binding sites. During this process, we extracted three features - motif structure, motif sequence, and motif set - and conducted a quantitative analysis. The study of Daniel *et al*. on binding preference of human RBPs revealed that RBPs have a tendency to bind RNA motifs with low complexity and large number of motifs with strong binding preferences were abundant in the single-base-rich and A/U-rich or C/U-rich regions (Dominguez et al. 2018). Based on this finding, we calculated the Shannon entropy of each motif in the sliding window and defined motifs with a Shannon entropy less than 1.1 as binding preference motifs (Figure 1C). Subsequently, we constructed Position Probability Matrices (PPMs) based on the secondary structure and sequence of these selected binding preference motifs. To facilitate PPM description, we categorized secondary structure features into five units: B (Bulge), E (External strand), H (Hair loop), M (Multiple loop), and S (Stem). Furthermore, for each selected binding preference motif, it was named after the base with the highest count to generate a set of all motifs (Figure 1C). Pearson correlation coefficients are computed for these three features separately at both native and predicted binding sites, and subsequently integrated into LS-seq to assess the similarity in sequence binding preferences.

### Optimizing the scoring strategy based on LS-PEAK

The test revealed that utilizing only LS-Score as a filter for the templates (Top5) resulted in an average MCC of approximately 0.25 for the predicted outcomes. To investigate the underlying reasons and take statistical information of LS-Score into account, the following procedures were implemented for each all-to-all alignment. The goal was to identify 8-mer RNA motifs on the target sequence that exhibit strong binding preferences and significant sequence similarity with native binding sites. Therefore, we took into account the LS-Scores of all results of all-to-all alignment, instead of only taking the Top5.

First, LS-Scores are evenly distributed to the bases (*v*) of motifs (*N*_moti*f*s_, number of the 8-mer motifs) screened by low Shannon entropy algorithm at the predicted binding sites:

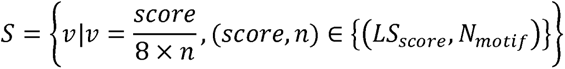

Then *v* is accumulated to calculate the *Value_i_* of each *base_i_* on the target RNA sequence:

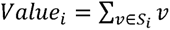

Utilizing these Values, we can generate the binding preference peak map of the target RNA at base-resolution, hereafter referred to as LS-PEAK.

In LS-PEAK, the highest five peaks are selected as potential strong binding preference regions. Then, the top 100 predicted binding sites ranked by LS-Score are selected, and the distance from their midpoint to these five peaks is calculated. Finally, the five predicted binding sites closest to the five peaks are selected as the output of the results. The rationale for this choice is discussed in the RESULTS section.

## RESULTS

We present PRIME-BSPre, a genome-wide protein-RNA binding sites prediction method based on templates. Its alignment and template screening process incorporates both the RNA sequences, secondary structures, and the corresponding tertiary structures of RBPs. Compared with some deep learning methods, we transcend the constraints of one model for one protein and exhibit robust predictive power for RBPs beyond the template library and in different cell lines. In a comparison with PrismNet we found that when their model trained in HepG2 was used to predict the same RBPs in HEK293, the prediction performance dropped significantly. We speculate that this may be due to the fact that during training, the model learns some specific features caused by cell lines, experimental methods and the environment in addition to binding preferences, which may further reduce the universality of the model. However, RRIME-BSPre can still achieve an impressive prediction success rate in different cell lines (Table1). The LS-PEAK metric, provided by PRIME-BSPre, not only enhances the efficacy of template screening but also enables direct prediction of RNA binding preference motifs on target sequences due to its composition (LS-Score), which primarily reflects the sequence-specific binding preferences of RNAs. We compared LS-PEAK with peak plots generated by Z-Score (Z-PEAK for short) calculated by RBPmap (all Human/Mouse motifs mode) (Paz et al. 2014), and the average Pearson correlation coefficient reached about 0.6. This observation suggests a strong correlation between the LS-PEAK, obtained through screening 8-mer motifs with low Shannon entropy, and the Z-PEAK derived from native RNA-binding motifs as templates.

### Correlation analysis between scores and MCC

We analyzed the RNAs in each RBP data set and compared the correlation between the scores (LS-Score, LS-seq, LS-2d) obtained by each template and its predicted MCC using Pearson correlation coefficient (Figure 2A). Among the boxplots, LS-seq shows the strongest correlation with MCC (PCC=0.30). As LS-Score is primarily determined by LS-seq due to the scoring strategy (Table S2), their correlation distributions are similar (PCC=0.28). In contrast, except for a few outliers, there is a weak correlation between LS-2d and MCC (PCC=0.04). It suggests that sequence binding preference is more prevalent than structure binding preference. Besides, the LS-2d outliers do provide us with some templates that exhibit high MCC in cases where sequence binding preference is weak. For instance, ZNF75A_ENST00000612186.1 (binds to ZC3H7B) and its Top1 template ATP5IF1_ENST00000497986.5 (binds to HNRNPK) exhibited strong similarity in secondary structure (LS-2d=0.853), but low similarity in sequence binding preference (LS-seq=0). Moreover, due to the intricate nature of protein-RNA interaction, which still harbors numerous ambiguous details and unestablished rules, our scoring function alone can merely accomplish preliminary template screening, exhibiting only a moderate correlation with MCC. To further enhance the accuracy, it is imperative to integrate both LS-Score and LS-PEAK as previously mentioned.

**Figure 2.**
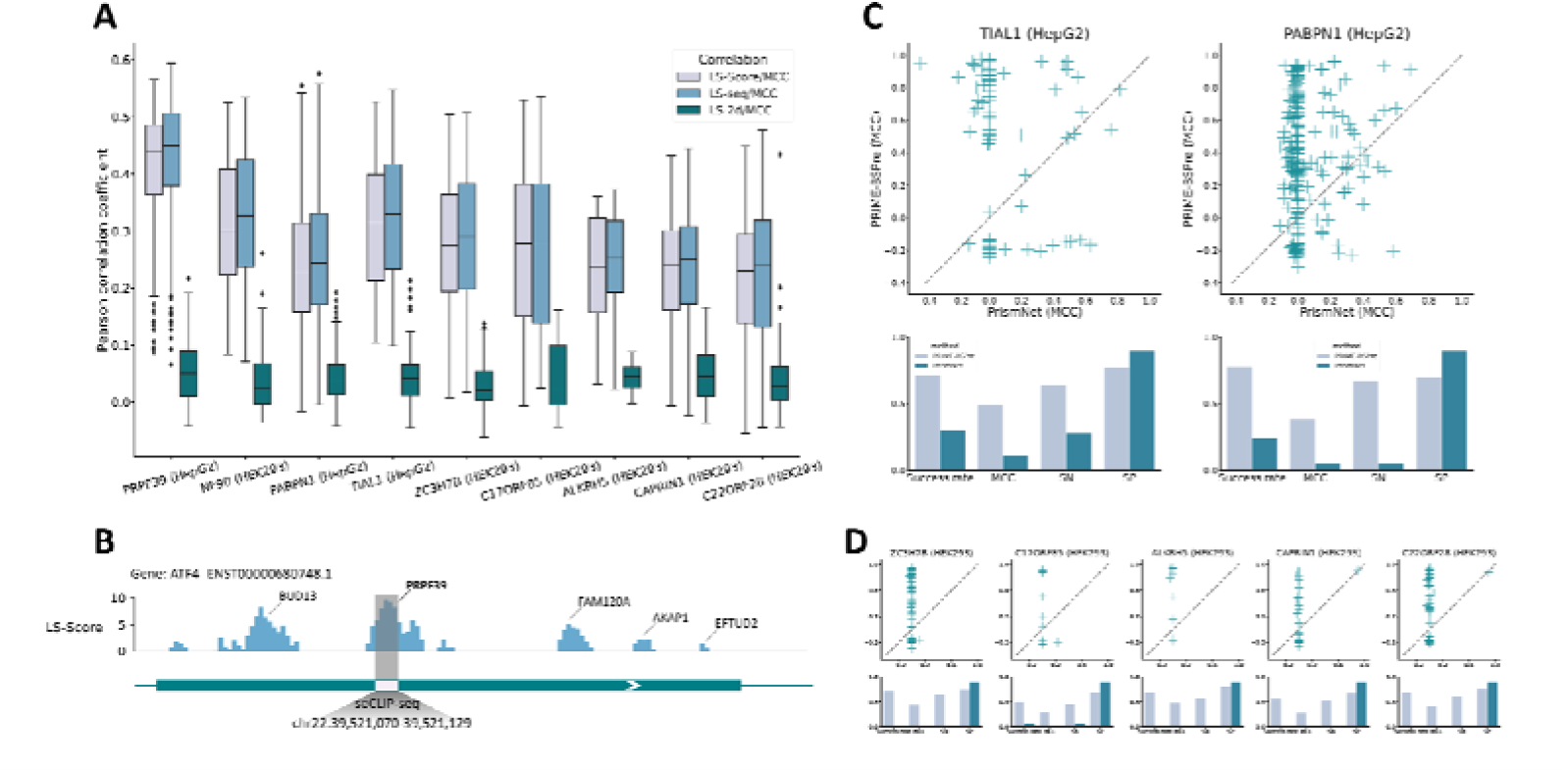
Analysis of scoring and comparison with PrismNet on different cell lines. **A** Correlation between template screening scores and MCC. Each target RNA undergoes an all-to-all aligned with the 15,348 RNAs in the template library. By calculating the Pearson correlation coefficient between the MCC predicted by all templates and their corresponding scores (LS-Score, LS-seq, LS-2d), we analyzed the effectiveness of these scores and obtained insights into the importance of their respective features in the binding process. **B** Peak plot given by PRIME-BSPre for regions with binding preferences on ATF4_ENST00000680748.1 (LS-PEAK obtained from seCLIP-Esq. **C-D** Comparison of PRIME-BSPre with PrismNet in different cell lines. Dot plots depict the performance of all RNAs in each RBP, as predicted by PRIME-BSPre and PrismNet.

#### LS-PEAK and its biological interpretation

In the Method section, we have provided a detailed explanation of how to generate LS-PEAK and utilize it for optimizing template screening. To summarize, LS-PEAK is a statistical peak map under base-resolution. We hypothesize that the regions where peaks are located contain higher binding preferences and are more likely to be potential binding sites. Occasionally, LS-PEAK may exhibit multiple peaks, and our study revealed that some of these peaks, aside from the Top1 peak, should not be considered as noise. In the eCLIP data, we observed not only the general phenomenon of different binding sites being associated with distinct RBPs but also numerous instances where multiple regions on a single transcript bound to the same RBP and different RBPs interacted with the same location on one transcript. In the process of analyzing LS-PEAK, we have also observed that certain sub-peaks exhibited interactions with other RNA-binding proteins (RBPs) within the template library, which aligns with the findings obtained from eCLIP data. Taking the prediction of binding sites on ATF4_ENST00000680748.1 interacting with the protein PRPF39 as an example (Figure 2B), the mean value of all LS-Scores was used as the threshold to intercept LS-PEAK, and some separate peaks were kept. The main peak was experimentally demonstrated to bind to PRPF39, and the sub-peaks were also experimentally proven to be native binding sites that interact with other proteins. However, in prediction, it is not clear whether there is only one binding site on the target RNA corresponding to the target RBP. Therefore, we choose the Top 5 peaks as the candidate regions and select the five templates closest to these peaks as our final output results.

### Compare PRIME-BSPre with PrismNet to demonstrate its applicability across diverse RBPs and cell lines

PrismNet(Xu et al. 2023), a newly launched server for predicting protein-RNA binding sites, outperforms other mainstream deep learning methods and also incorporates RNA sequence and secondary structure characteristics. The PrismNet outputs we used here were derived from the “Pre-calculated RBP binding sites” provided by its web server. To ensure objectivity in our comparison, we selected CLIP data on HepG2(Xu et al. 2023) and HEK293(Baltz et al. 2012) for testing, as a significant proportion of the templates in PRIME-BSPre library were derived from the K562. While about PrismNet, we chose their sequence & structure models trained on HepG2. TIAL1 dataset (containing 76 RNAs) and PABPN1 dataset (containing 238 RNAs) were chosen as the two RBPs for comparison on HepG2 (Figure 2C). The results showed that PRRIM-BSPre performed better, with a prediction success rate of 71% [54/76] on TIAL1 (defined as success if MCC over 0.1), averaging MCC=0.50, and a success rate of 78% [185/238] on PABPN1, averaging MCC=0.38. In contrast, the success rate of PrismNet was only 30% [23/76] on TIAL1, with an average MCC=0.10, and that of PABPN1 was 21% [51/238], with an average MCC=0.05. It is noteworthy that the high specificity observed in both methods (Figure 2C-D) can be attributed to the significantly shorter predicted binding sites compared to the full-length of the RNAs. This results in a high true negative rate for all predictions, so the validation of prediction requires both SN and SP to reach a high value simultaneously. Subsequently, we utilized RNA binding sites data from five RBPs (ZC3H7B, C17ORF85, ALKBH5, CAPRIN1, C22ORF28) obtained through PAR-CLIP experiments on HEK293 conducted by Baltz *et al*. to compare the predictive performance of the two methods across different cell lines. The PrismNet’s model used to predict was also trained on HepG2 and PRIME-BSPre still relied on the original template library. Our results demonstrate that while PrismNet models trained on HepG2 exhibit limited capability in predicting these RBPs, PRIME-BSPre consistently maintains its robust predictive power.

The utilization of deep learning methods for predicting RNA binding sites, as exemplified by PrismNet, still faces numerous challenges. Firstly, each RBP requires a distinct model to be trained, which consequently necessitates data on protein-RNA interactions. This significantly raises the threshold as predictions are often made solely due to insufficient experimental data. Additionally, a model trained on one cell line found it difficult to transfer to other cell lines in the aforementioned comparison. The model seems to have acquired distinct binding preferences for each cell line or may have been influenced by the experimental conditions. In summary, although models trained using deep learning methods can accomplish some prediction tasks, they are far less stable and portable than template-based methods.

### The comparison with RBPmap demonstrates the effectiveness of low Shannon entropy feature in identifying RNA motifs with strong binding preference

RBPmap predicts RNA binding sites using an RNA binding motif database. By applying its Weight-Rank algorithm(Akerman et al. 2009), RBPmap maps a given RNA motif to the most appropriate position on the target RNA sequence, which is then predicted as the binding sites. If we want to predict the interaction between an RBP and an RNA using RBPmap, it is crucial to have knowledge about the specific RNA motifs preferred by the RBP. In cases where this information is missing from the database provided by RBPmap, prediction becomes challenging. However, if the goal is solely to determine the binding preference region on the target RNA without specifying any particular RBPs, our select the “All Human/Mouse motifs” in the “Motif selection” section of the server (http://rbpmap.technion.ac.il/) for computation. Then the Z-Scores of all identified RNA motifs are allocated into their respective base positions, ultimately generating a peak map resembling LS-PEAK (referred to as Z- PEAK hereafter). Daniel *et al*. have mentioned in their studies that RBPs prefer to bind low-complexity motifs, and this preference has in turn led to the convergence of preferences among various RBPs in evolution (Dominguez et al. 2018). Therefore, upon traversing the motifs, if certain regions exhibit elevated scores (representing Z- PEAK peak regions), the inference can be made that these motifs may exhibit a concurrent binding preference for multiple RNA-binding proteins (RBPs). Consequently, it is reasonable to classify them as potential binding sites due to their shared binding characteristics.

The above is an innovative application of RBPmap, which is based on the construction of LS-PEAK, wherein it is important to note that while Z-PEAK and LS-PEAK share similar generation logic by assigning scores to corresponding base regions on the RNA sequence, Z-PEAK is generated based on an existing RNA binding motif database, whereas LS-PEAK is obtained through sequence similarity alignment and low complexity screening. We assessed the correlation between LS-PEAK and Z-PEAK as well as the overlap rate of their Top5 peaks, revealing a strong correlation between these two peak plots (evaluated across all 665 RNAs; mean PCC=0.55; Top5 peaks overlap rate = 77%) (Figure 3B). Based on this, we employed Z-PEAK in the template screening process following all-to-all alignments and compared its predicted performance with that achieved using LS-Score ranking alongside LS-PEAK optimization (Figure 3A). The effective prediction region of both screening methods is located in the upper right corner of the two-dimensional KDE (kernel density estimation) plot. It is evident that employing LS-Score and LS-PEAK generally outperforms using Z-PEAK, as the high-density area is concentrated above the dashed line. Nevertheless, our screening algorithm still demonstrates good performance in regions where Z-PEAK fails to predict, specifically in the top left box.

**Figure 3.**
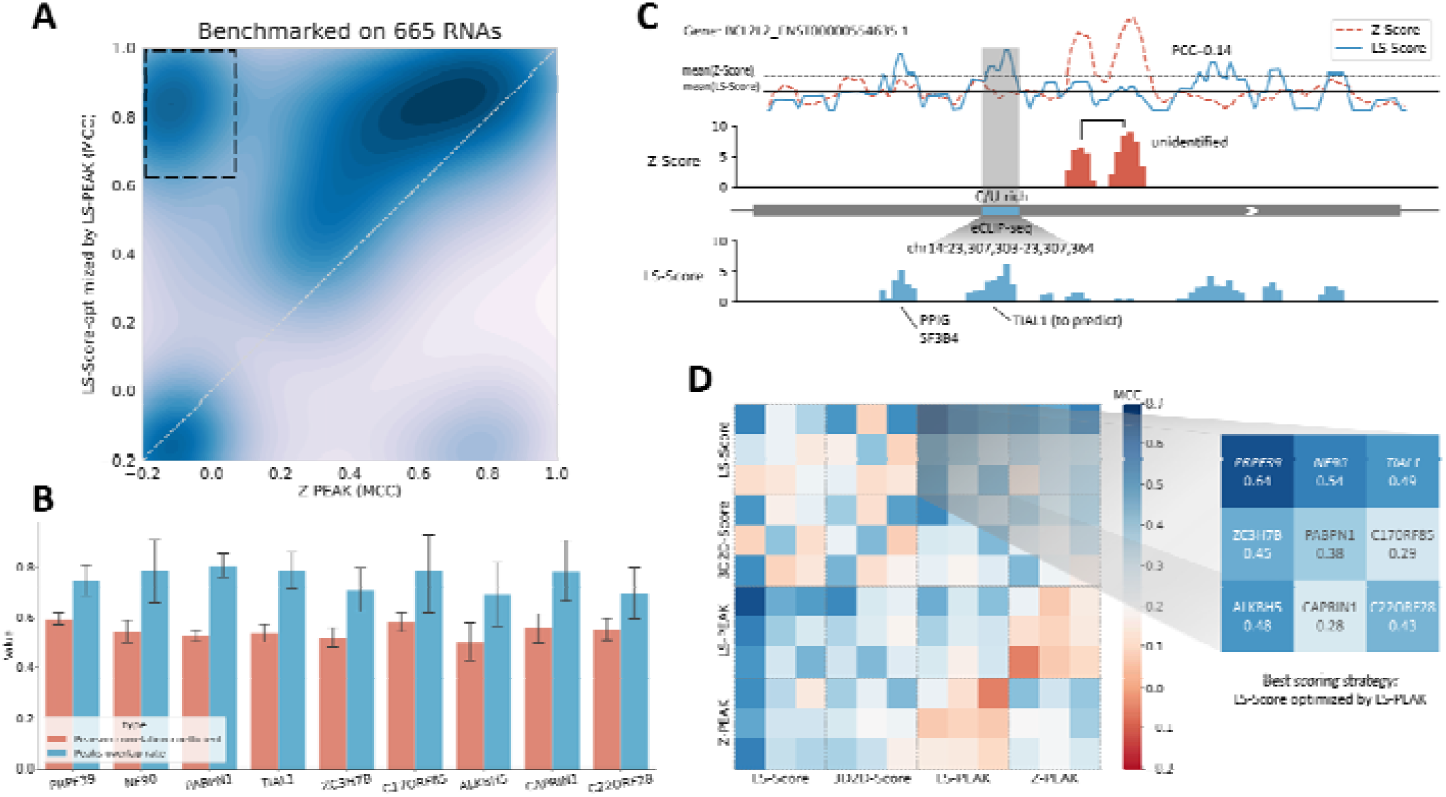
Comparison with RBPmap and comparative analysis of all scoring strategies. **A** Compare the effect of PRIME-BSPre scoring strategy and Z-PEAK on screening template. The dark hues indicate the area of high density in the two-dimensional kernel density estimation. **B** Pearson correlation coefficient of LS-PEAK and Z-PEAK on different RBP datasets and the overlap rate of their respective Top5 peaks. **C** A comparative analysis of LS-PEAK and Z-PEAK, using BCL2L2_ENST00000554635.1 as an example. **D** Heat-map comparison of all the scoring strategies. Each dashed square represents a scoring strategy, with each color block within the square indicating the predicted outcome of an RBP dataset.

Z-PEAK was able to identify some positive predictions from all-to-all alignment, resulting in even better performance than PrismNet. This demonstrates the effectiveness of our innovative integration of RBPmap output (z-score). However, direct screening with Z-PEAK has limitations. Firstly, it overlooks secondary structure and sequence similarity within the contexts of RNA motifs which have been found relevant for protein-RNA interactions (Fu and Ares 2014; Taliaferro et al. 2016; Dominguez et al. 2018). Secondly, although the peaks of LS-PEAK and Z-PEAK overlap by 77%, differences in prediction performance may arise from some non-overlapping peaks. For instance, taking BCL2L2_ENST00000554635.1 as an example (Figure 3C), the peak maps show that the correlation between LS-PEAK and Z-PEAK is weak, and there is divergence in the binding sites. After investigation, it was discovered that the binding sites of this RNA exhibited a high frequency of C/U-rich motifs. But only 7 out of the 233 motifs (corresponding to 132 human/mouse RBPs) in the RBPmap database were found to be C/U rich. Therefore, without considering the distribution of various characteristic motifs, Z-PEAK may not effectively identify binding sites that contain these rare motifs. In contrast, our template library includes a substantial number (15348) of RNA templates for 158 human RBPs, which partially addresses this limitation. Moreover, among the Top100 templates ranked by LS-Score, selection is not solely based on local binding preferences but also considers the high similarity in secondary structure and sequence on a larger scale which takes the contexts into consideration. This richer information combined with subsequent LS-PEAK optimization enhances confidence and accuracy in prediction.

### Benchmark PRIME-BSPre on different cell lines and conduct a comprehensive comparison of all scoring strategies

We benchmarked PRIME-BSPre on two cell lines, including two recent CLIP datasets, PRPF39 (on HepG2) obtained through seCLIP-seq by Blue et al. and NF90 (on HEK293) obtained through iCLIP by Lodde et al. (Blue et al. 2022; Lodde et al. 2022). The results achieved in comparison with PRIME-3D2D and PrismNet are remarkably superior (since PrismNet did not train models for these two RBPs, they were excluded in the PrismNet evaluation) (Table1).

Additionally, we conducted a systematic analysis of various scoring strategies utilized in the research and their combinations (Figure 3D). Our findings indicate that implementing LS-PEAK optimization on the Top100 templates ranked by LS-Score yielded the best predictive performance, resulting in a 76% increase in MCC compared to utilizing LS-Score ranking alone. Notably, our prediction performance on two experimental datasets PRPF39 (MCC=0.64) and NF90 (MCC=0.54) was striking.

## DISCUSSION

PRIME-BSPre is a computational method that predicts RNA binding sites through all-to-all alignment, taking into account sequence similarity, secondary structure similarity, and motifs binding preference. Compared to current deep learning methods, PRIME-BSPre overcomes the challenge of requiring experimental data for training before prediction and exhibits greater robustness in predicting performance across different cell lines. Our benchmarks were conducted on complete transcripts rather than artificial fragments flanked by specific lengths. Additionally, we are the first to introduce a low Shannon entropy algorithm to describe motif binding preferences. The predicted results also support the notion that RBPs are more likely to bind low-complexity RNA motifs.

Given the genome-wide nature of our prediction, we have observed that there exist some binding sites located within repetitive genomic regions, which are prevalent throughout the human genome (de Koning et al. 2011; Gemmell 2021). Notably, some binding sites appear to be closely associated with these repetitive regions, while others occur between repeat areas or even coincide with them. Moreover, these repetitive regions also exhibit characteristics of low complexity, such as continuous repetition of double bases or repeats in the form of a triple or quadruple that initiates with a single base. Some templates in our library contain binding sites within the repeat region, which are represented in lowercase letters in the genome. The scoring of sequence similarity takes the capitalization of bases into consideration, thereby enabling the prediction of certain binding sites based on this feature. NF90, previously mentioned, exhibits multiple repetitive regions within its interacting RNA transcripts, some of which encompass binding sites. Investigating the intricate interplay between these repeat regions and RBPs holds substantial scientific significance.

## DATA ACCESS

All the comparison datasets and their prediction performances have been uploaded at http://rnabinding.com/dataset_performance.rar.

Code is available under http://rnabinding.com/PRIME-BSPre.tar.gz.

## ACKNOWLEDGEMENTS

The computation is completed in the HPC Platform of Huazhong University of Science and Technology. This work has been supported by the National Natural Science Foundation of China [32271267].

